# Predictive analysis of patient recovery from cardiac-respiratory arrest

**DOI:** 10.1101/650408

**Authors:** A. Floyrac, A. Doumergue, N. Kubis, D. Holcman

## Abstract

The severity of neuronal damages in comatose patients following anoxic brain injury can be probed by evoked auditory responses. However, it remains challenging to predict the return to full consciousness of post-anoxic coma of hospitalized patients. We presented here a method to predict the return to consciousness based on the analysis of periodic responses to auditory stimulations, recorded from surface cranial electrodes. The input data are event-related potentials (ERPs), recorded non-invasively with electro-encephalography (EEG). We extracted several novel features from the time series responses in a window of few hundreds of milliseconds from deviant and non-deviant auditory stimulations. We use these features to construct two-dimensional statistical maps, that show two separated clusters for recovered (conscience) and deceased patients, leading to a high classification success as tested by a cross-validation procedure. Finally, using Gaussian, K-neighborhood and SVM classifiers, we construct probabilistic maps to predict the outcome of post-anoxic coma. To conclude, statistics of deviant and non-deviant responses considered separately provide complementary and confirmatory predictions for the outcome of anoxic coma.

## 1 Introduction

Sudden death by cardiac arrest is a major public health problem, affecting 55 patients out of 100,000 with nearly 40,000 cases per year in France. When cardiac arrest happens in a hospital, the chances of survival at one year are about 30% but drops to 5% when cardiac arrest occurs outside a hospital. Among the survivors, 68% will have a moderate disability in the following year. To limit brain damage, patients with post-anoxic coma are now exposed to therapeutic hypothermia for 24 hours. An evaluation of the neurological damage is ideally performed 48 hours after the interruption of the sedative treatment to determine the neurological prognosis. It is a multimodal evaluation combining a clinical evaluation (Glasgow Scale, photomotor and pupillary reflexes) and electrophysiology, an evaluation of biological markers that can complete this series of examinations (NSE and S100b proteins to determine neuronal necrosis). Post-anoxic encephalopathy and its prognosis is also evaluated by electroencephalogram (EEG) [1]: the absence of a N20 response to somatosensory evoked potentials after stimulation of the median nerve has a specificity almost equal to 100% to predict the absence of awakening in the adult. However, the lack of response in intensive care unit (ICU) patients is difficult to assert due to the electrical environment that generates many artifacts that sometimes make it extremely difficult to interpret the low amplitude response of the evoked potential. In addition, the lack of response to this assessment predicts a poor prognosis.

In general, from electrodes placed on the scalp, evoked potential responses (EPR) are recorded, reflecting the brain response to auditory stimulation. From the EPR, the MisMatch Negativity (MMN) signal is computed [2] as a negative potential between the potentials produced by regular and deviant sounds. The difference between the two responses points is expected to occur between 150 and 300ms [3] and represents the cerebral integration of sensory memory in response to infrequent and random changes in a continuous time series of sounds. Indeed, standard sounds differ from deviant sounds in their intensity and frequencies. The study of MMN is performed by averaging responses to different stimuli to reduce the impact of brain activity independent of the stimulus [4, 2, 1]. However, the analysis and recognition of the presence or absence of MMN is difficult in intensive care patients, as many artifacts and a drastic reduction in the amplitude of evoked potentials make them difficult to isolate.

The results are not encouraging with the number of standard electrodes and resuscitation conditions, and the MMN prediction seems to have little positive and negative predictive value. Indeed, a considerable amount of information is lost by considering only these averages and calls for a different statistical approach for the diagnosis of the patient brain condition. Several complementary methods have been developed to quantify the predictive value of the MMN for comatose patients: some are based on t-test in time (100-200 ms) to check the presence of a detectable peak of the mismatch negativity, others are based on wavelet transform, multivariate, cross-correlation and probabilistic methods [4, 5, 6, 7, 8, 9, 10, 11]. Finally, we still missing a consensus about the quality of the existing features [12, 2, 13] to predict the outcome of post-anoxic coma.

We develop here an approach based on the statistical identification and classification of transient features, present in the EEG responses and a probabilistic classifier to predict the outcome of post-anoxic coma for hospitalized patients. The input data are event-related potentials (ERPs), recorded non-invasively with electro-encephalography (EEG). We change from the classical paradigm and study separately deviant and non-deviant responses using the new identified patterns, that are different from the mismatch negativity. For deviant responses, one novel feature is the local oscillation of the ERP signal. Moreover, we estimated the probability for comatose patients to return to consciousness. We use several classifiers that converge in predicting the outcome of post anoxic coma. Finally, using deviant and nondeviant responses, our probabilistic classifiers allow to represent the empirical data into a two-dimensional map that we construct. These various maps can be upgraded when adding a new classified case, a procedure that increases the overall performance of the present probabilistic method.

## 2 Results

### 2.1 Pre-processing of deviant and non-deviant responses

To quantify the auditory evoked responses recorded from post-anoxic coma patient in intensive care, we focus on the non-deviant and deviant stimulation (Material and Method), stimulated every seconds. The signal was recorded from various electrodes (Fig. 1A) and there was one deviant for ten non-deviant sounds. We study each response to standard and to deviant stimulation separately (Fig. 1B-C and SI Fig. S1). For each of the response, we use the Cz-electrode only for the non-deviant stimulation and average over the main electrodes for the deviant response and then filtered the signal in a band [0.5-50]Hz (material and methods). Finally, we average over all stimulations, leading to a response in the time interval [0 − 1000]*ms* (non-deviant) and [0 − 500*ms*] (deviant). To classify each response separately, we identity different features that could characterize the responses in each cases (Fig. 2 and Fig. 3).

**Figure 1:**
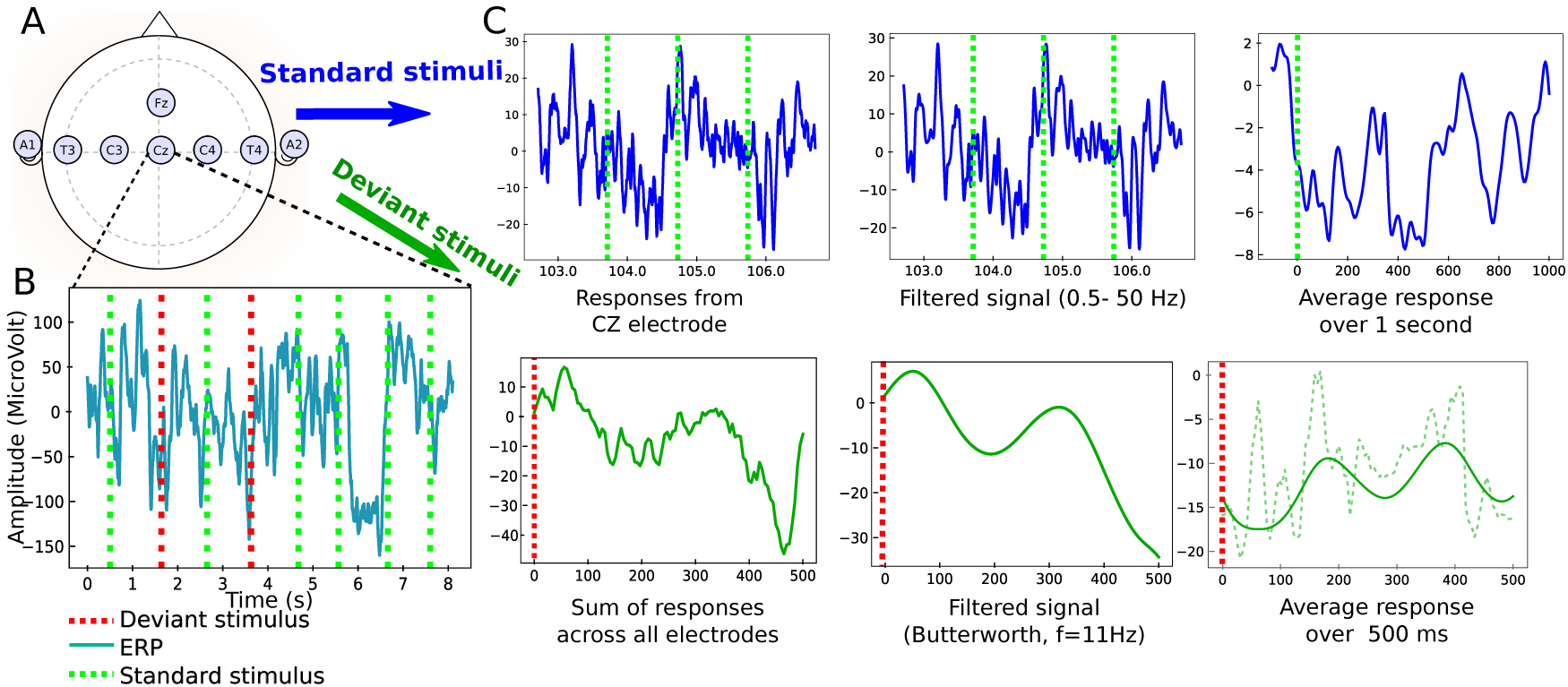
Pre-processing of the Evoked Auditory Responses to standard and deviant stimuli. **A.** Upper: standard position of the EEG electrodes. Lower: EEG traces during a protocole mixing deviant (red) and non-deviant (green) stimulations. **B.**: sample of standard stimuli (blue) the EEG signal from CZ-electrode is filtered 0.5-50Hz. The output is an average filtered response over one second. **C.** Pre-processing of deviant stimuli: 1) the signal is summed over electrodes, 2) a low-pass filter is applied (Butterworth with *n* = 2, cutoff frequency at 10Hz). 3)Average filtered response (continuous green) in a window of 500*ms* to a deviant stimulus, computed after synchronization to the stimulus. The non filtered averaged response is also shown (dashed line).

**Figure 2:**
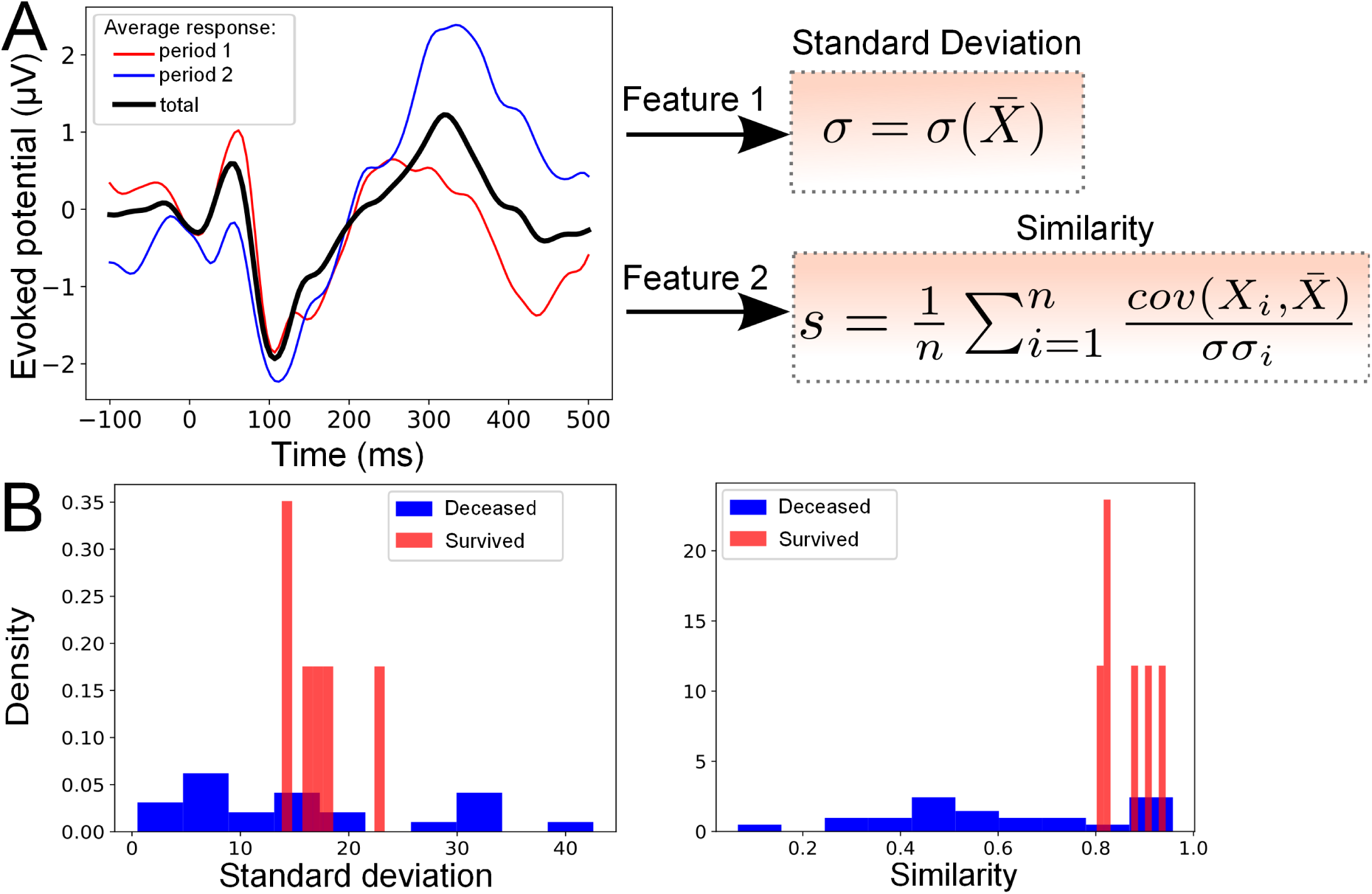
Statistical features associated to non-deviant responses. **A.**Left: Average evoked responses computed over a time window of [0 − 10]*min* (period 1,red), [10 − 20]*min* (period 2,blue) and over the entire period ([0 − 20]*min*, black). Right: The standard deviation *σ* and the average correlation function *s* (Similarity), between the response over the entire period ([0−20]*min*) and over one of the n periods ([0−20]*min* or [10−20]*min*). Here *n* = 2. **B.** Feature distributions. Distribution of the standard deviation (Left) and similarity (Right), computed over the entire period and dataset of patients from the Cz-electrode: Red (survived) and Blue (deceased).

**Figure 3:**
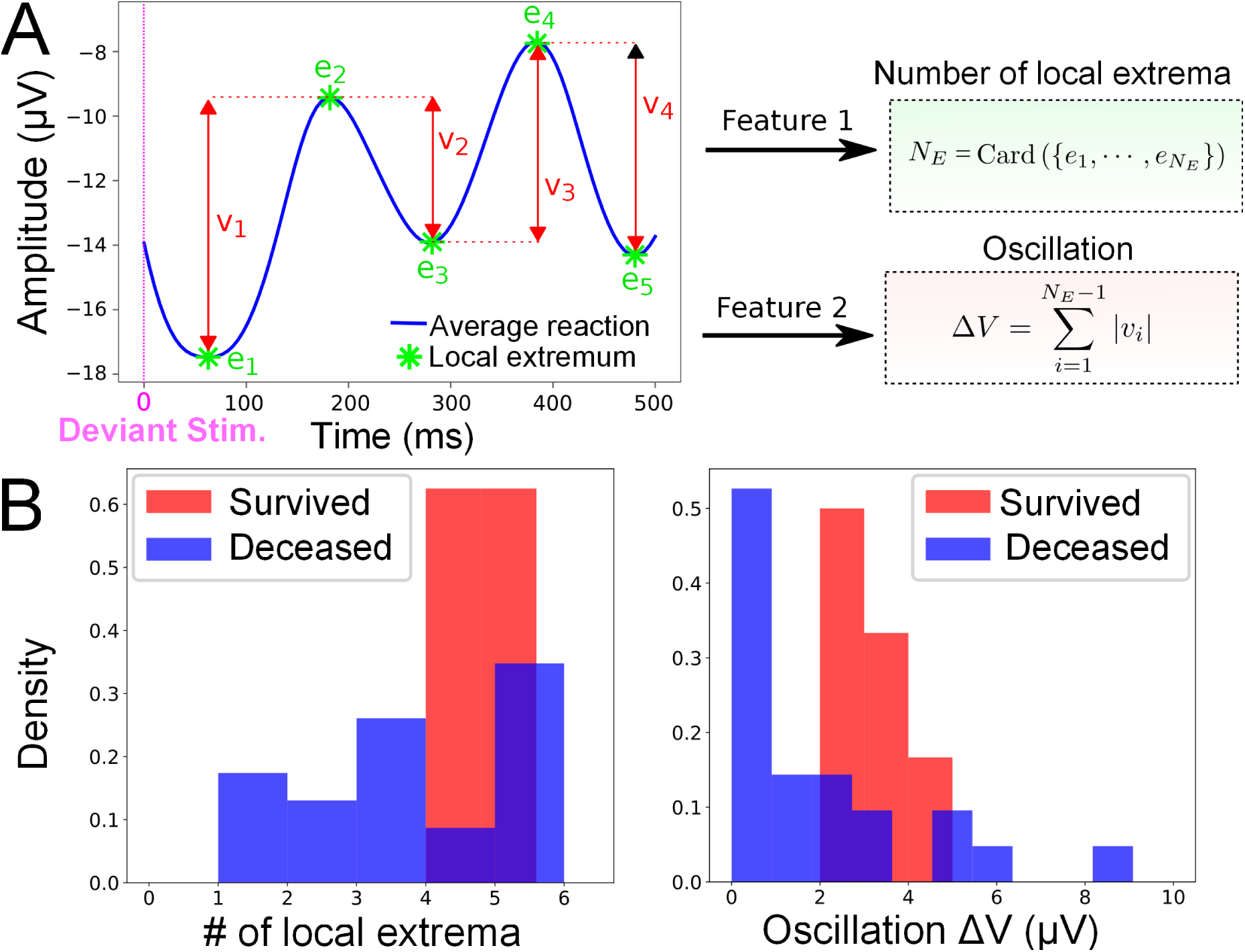
Statistical features associated to deviant responses. **A.**Left: The average filtered evoked response (blue) to deviant stimuli computed over the entire time window contains *N*_*E*_ local extrema *e*_*i*_ (minimum or maximum), which is the first feature. The second feature is the oscillation |Δ*V* | =∑_*i*_ |*V* (*e*_*i*_) − *V* (*e*_*i*+1_)|, which is the sum of the absolute value of the difference between two consecutive extrema of the average evoked response (see fig. 1C-step3). Feature distributions. Left: Distribution of the oscillation (Left) and the number of local extrema (Right) computed over the entire period and dataset of patients from all electrodes: Red (survived) and Blue (deceased).

### 2.2 Feature identification associated to deviant and non-deviant responses

For non-deviant responses, we subdivided the entire 20 minutes of recording into two time windows: the first 10 minutes (first period) and the period 10 − 20minutes (second period). We computed the variance (formula 5) and the correlation function (formula 6) of the response computed between the response in the first and second time period (Fig. 2A-Right). To test the ability of these two parameters to separate deceased from survival patients, we plotted the histogram of these two parameters for all patient in our data (Fig. 2B), showing that each individual feature could potentially be used for a robust classification (see below).

For the deviant responses, the signal showed different characteristics and we thus decided to use different features: the first one consists in the number of extremums *N*_*E*_ in the signal (Fig. 3A) and the second is the absolute value of the oscillation|Δ*V*|, which represents the sum of the differences between the extrema (formula 8). The result of the classification is shown by histograms of the two parameters computed over the whole population (Fig. 3B).

At this stage, we conclude that we selected two different types of parameters to study deviant and non-deviant responses, but each of them taken individually is not sufficient to clearly separate survivors from deceased patients.

### 2.3 Survival map constructed from Bayesian statistics

Since each parameter taken individually for deviant and non deviant responses are not sufficient to obtain a clear separation between the population of deceased and surviving patients, we decided to combine them into a two-dimensional map (Fig. 4). Interestingly, we found that this map allowed a clear separation and we quantified this separation using various a priori classifiers: for non-deviant responses, we use successively SVM, Gaussian and the K-neighbor classifiers[14]. The classification probability of a patient characterized by its coordinates is obtained by computing a score that measures the proximity to one of the two classes(Material and method section 4.6).

**Figure 4:**
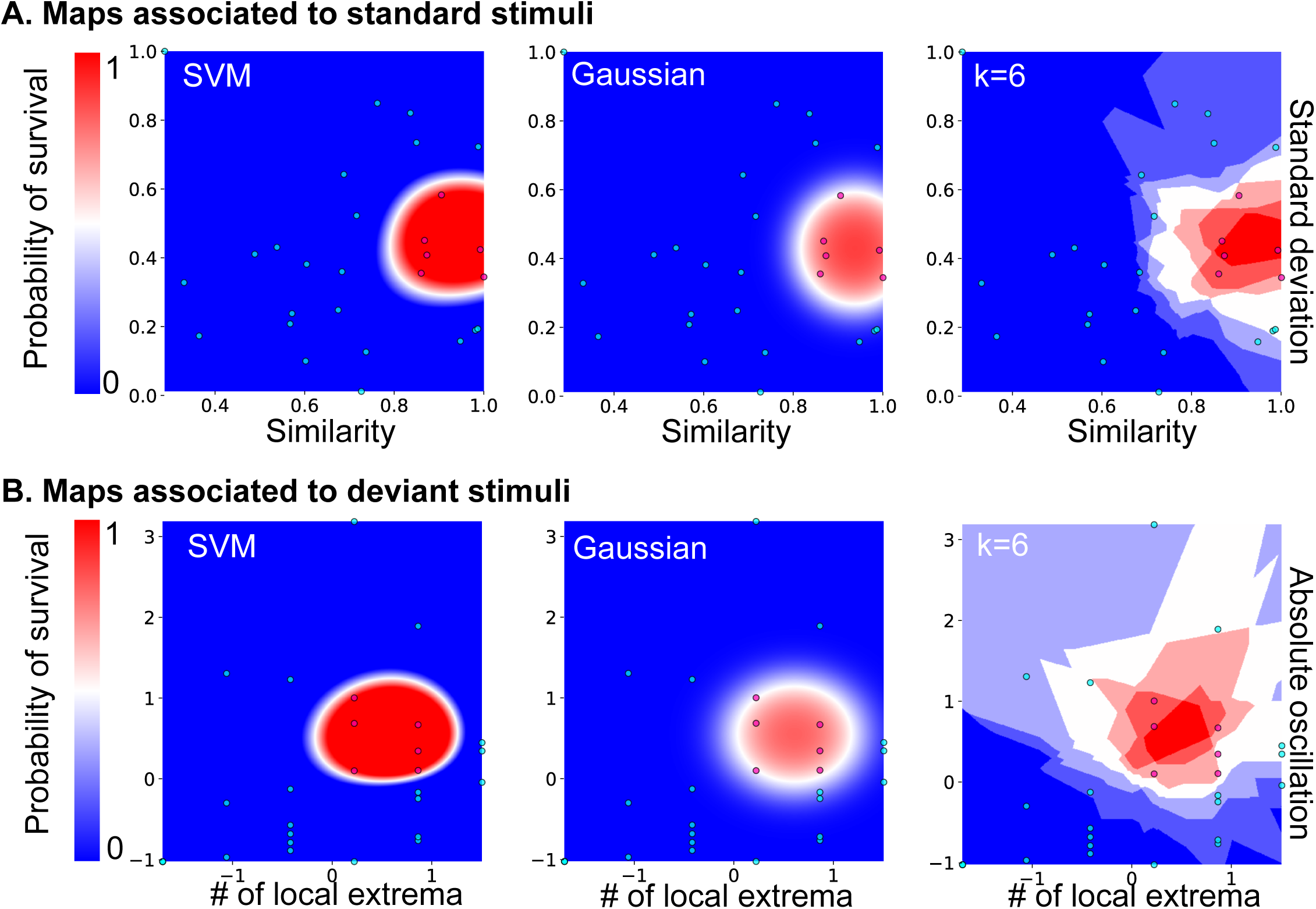
Predictive probability maps of survival. **A** Probability maps computed from features of the standard stimuli responses (fig. 2). From Left to right: maps computed from SVM, Gaussian and the k-Nearest Neighbors classifier (k=6). **B** Probability maps computed using deviant stimuli features (fig. 3). From Left to right: maps computed from SVM, Gaussian and the k-Nearest Neighbors classifier with distance-related weights for k=6.

Surprisingly, we found that survival patients aggregate into a bounded region(red) (Fig. 4A) and well separated from the region associated to deceased patients. This partition between two distinct regions was present for all classifiers: SVM, Gaussian and k-neighbors, confirming that this partition was robust independently of the choice of the classifiers (see SI Fig. S5, for other choices of *k* for the k-neighbor algorithm). Interestingly, in parallel we also tested our classification approach for deviant responses, using the associated features (number of extremum *N*_*E*_ and oscillation |Δ*V*|). We found a similar partition into two classes of deceased and survived patients when classifying the deviant responses with the same accuracy level in both maps (Fig. 4B). To conclude, the present classification maps for deviant and non-deviant response show that the output of post-anoxic coma can be predicted (see table 2). To obtain a more accurate classifier, we used a different version of the K-neighbors classifier, where the weight depends on the distance between the point to classify and the K-nearest neighbors (see formula 14). Finally, combining the probability computed in each map, we propose a decision probability which is the minimum of the two estimated probabilities.

### 2.4 Comparing two-dimensional maps with the Mismatch-Negativity classification

To test the predictive strength of the deviant and standard response classification, we compare it with the MMN obtained by subtracting the deviant from the standard response (SI Fig. S2) either from all data set or after randomization so that the total number of deviant and standard response are equal (SI Fig. S3). In both cases, we found that MMN had always a poor predictive value compared to the three classification approaches (SVM, Gaussian K-neighbors). We also confirmed this result using a cross-validation approach where we computed the confusion matrix 16 for the true positive *T*_*p*_ (number of patients that survived and were correctly classified), true negative *T*_*n*_ (number of patients that deceased and were correctly classified) and false positive *F*_*p*_(number of patients that survived and were incorrectly classified) and false negative *F*_*n*_ (number of patients that deceased and were incorrectly classified). The results are shown in table 2-4-3. For SVM,

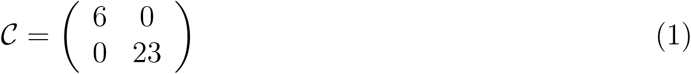

showing the classifier had 100% validation accuracy. We obtain such a quality for (*γ, C*) ∈ [0.5, 2.5] × [3, 30]. The confusion matrix computed for k-neighbors with distance-dependent weight is

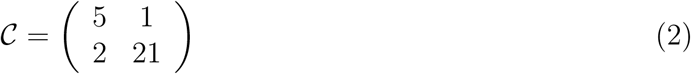

This estimator introduces type I error with an exactitude of 0.90, a sensitivity of 0.83 and a specificity 0.91. This estimator is less performant compared to SVM. The sensibility remains high and could be improved with the increasing number of classified patients. Finally, the confusion matrix obtained from MMN is

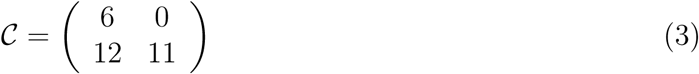

showing an exactitude of 0.59, a sensitivity of 1 and a specificity of 0.48. Thus confirming a MMN remains an interesting indicator for survival classification, but has a very weak specificity. All scores are summarize in table 1.

**Table 1:** Classification scores.

|  | Exactness | Sensitivity | Specifity |
| --- | --- | --- | --- |
| SVM | 1 | 1.0 | 1 |
| k-neighbors | 0.9 | 0.83 | 0.91 |
| MMN | 0.59 | 1 | 0.48 |

## 3 Discussion

We presented here an approach to predict from evoked auditory responses the output of anoxic coma. The method differs from previous ones which usually combine standard and deviant responses, that we considered separately. Average voltage topography analysis [9] already showed the difference in the two responses for control and coma patients. After we extracted here novel features from the time average responses, computed during the first 500*ms*, we constructed predictive maps in two dimensions. To evaluate the robustness of our method, we finally used three classifiers, showing similar maps classification results. Interestingly, the region of survival patients forms a bounded cluster, demonstrating that the EEGs of these patients have comparable and similar features. Finally, using cross-validation, we computed several classification scores, demonstrating that any of the three classification methods is more robust than the Mismatch negativity using logistic regression or single-trial topographic analysis [9]. At this stage, we conclude that survival probability maps allows us to predict the output of anoxic coma with a very good accuracy, sensitivity and specificity (tables 2-3). This approach that requires only a single electrode predicts and quantifies the survival rate based on a cohort but the prediction will improve as the number of classified patients increases.

**Table 2:**
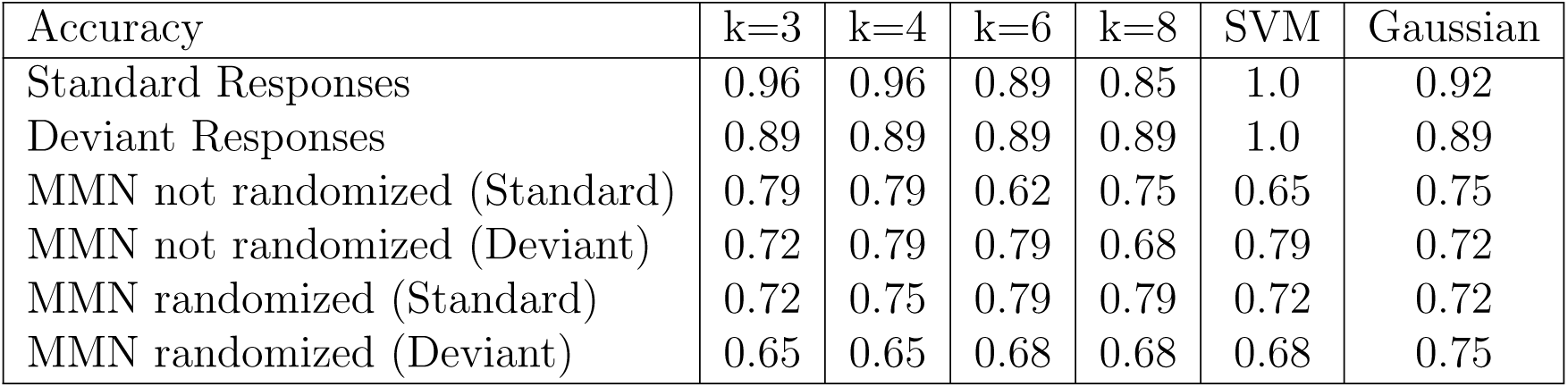
Accuracy rate obtained by cross-validation for responses to standard, deviant stimuli and MMN responses analyzed with the parameters of the standard and deviant stimuli.

| Accuracy | k=3 | k=4 | k=6 | k=8 | SVM | Gaussian |
| --- | --- | --- | --- | --- | --- | --- |
| Standard Responses | 0.96 | 0.96 | 0.89 | 0.85 | 1.0 | 0.92 |
| Deviant Responses | 0.89 | 0.89 | 0.89 | 0.89 | 1.0 | 0.89 |
| MMN not randomized (Standard) | 0.79 | 0.79 | 0.62 | 0.75 | 0.65 | 0.75 |
| MMN not randomized (Deviant) | 0.72 | 0.79 | 0.79 | 0.68 | 0.79 | 0.72 |
| MMN randomized (Standard) | 0.72 | 0.75 | 0.79 | 0.79 | 0.72 | 0.72 |
| MMN randomized (Deviant) | 0.65 | 0.65 | 0.68 | 0.68 | 0.68 | 0.75 |

**Table 3:**
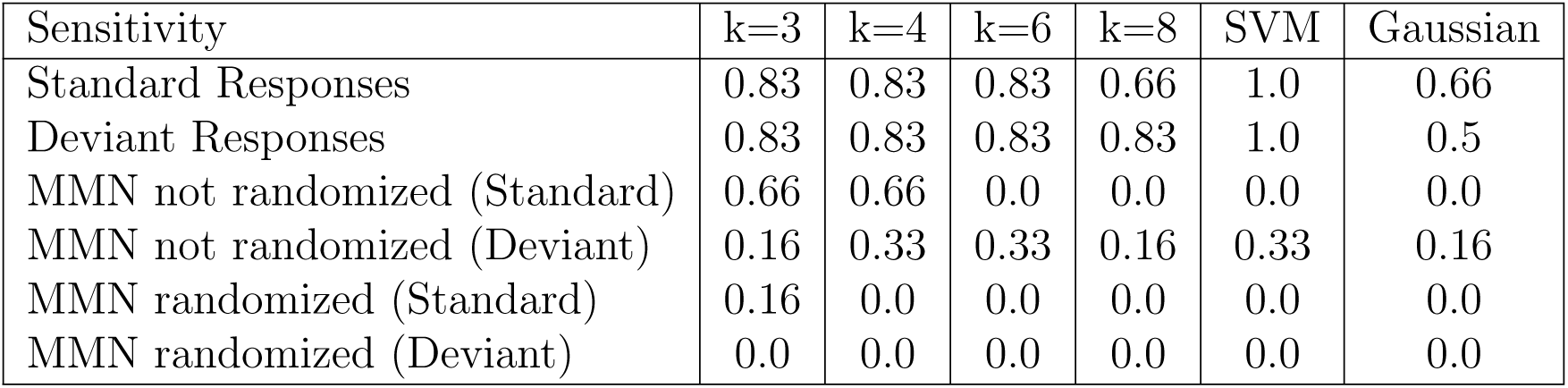
Sensitivity rate obtained by cross-validation for responses to standard, deviant stimuli and MMN responses analyzed with the parameters of the standard or deviant stimuli.

| Sensitivity | k=3 | k=4 | k=6 | k=8 | SVM | Gaussian |
| --- | --- | --- | --- | --- | --- | --- |
| Standard Responses | 0.83 | 0.83 | 0.83 | 0.66 | 1.0 | 0.66 |
| Deviant Responses | 0.83 | 0.83 | 0.83 | 0.83 | 1.0 | 0.5 |
| MMN not randomized (Standard) | 0.66 | 0.66 | 0.0 | 0.0 | 0.0 | 0.0 |
| MMN not randomized (Deviant) | 0.16 | 0.33 | 0.33 | 0.16 | 0.33 | 0.16 |
| MMN randomized (Standard) | 0.16 | 0.0 | 0.0 | 0.0 | 0.0 | 0.0 |
| MMN randomized (Deviant) | 0.0 | 0.0 | 0.0 | 0.0 | 0.0 | 0.0 |

**Table 4:** Specificity rate obtained by cross-validation for responses to standard, deviant stimuli and MMN responses analyzed with the parameters of the standard and deviant stimuli.

| Specificity | k=3 | k=4 | k=6 | k=8 | SVM | Gaussian |
| --- | --- | --- | --- | --- | --- | --- |
| Standard Responses | 1.0 | 1.0 | 0.9 | 0.9 | 1.0 | 1.0 |
| Deviant Responses | 0.91 | 0.91 | 0.91 | 0.91 | 1.0 | 1.0 |
| MMN not randomized (Standard) | 0.82 | 0.82 | 0.78 | 0.95 | 0.82 | 0.95 |
| MMN not randomized (Deviant) | 0.86 | 0.91 | 0.91 | 0.82 | 0.91 | 0.86 |
| MMN randomized (Standard) | 0.86 | 0.95 | 1.0 | 1.0 | 0.91 | 0.91 |
| MMN randomized (Deviant) | 0.82 | 0.82 | 0.86 | 0.86 | 0.86 | 0.95 |

### 3.1 Integrative value of the new features

Instead of using the MMN feature occurring in the range 150-300ms, we used here the cumulative oscillation of the ERP (Δ*V* relation 8) in the range of 20-500ms after stimulation. This analysis of the evoked response integrates various individual amplitude responses such as N20 to P300, which result from reverberation of neuronal networks activity in the frontal and auditory cortex [1]. In particular, the persistent presence of the P300 wave and large fluctuations of coherent response reflects a coherent brain activity. Conversely, the absence of such response or of any fluctuations is interpreted as significant damages. To conclude the oscillation feature defined in Fig. 3 and used in Fig. 4 reflects well the cumulative response of the Brain to an evoked responses.

The presence of the MMN was previously used to predict the outcome of anoxic coma [4, 7]. However, the MMN is not a specific marker because it is present in some comatose patients only [4] but is still used to predict a good outcome [15]. The probability maps reconstructed here from the MMN are much less fuzzy than the ones we obtained with the ERP responses (SI Fig. S4), with a much less predictive value and we decided not to use them. Recently, MMN was observed in sedated critically ill patients and participate in predicting awakening, but is again not robust enough to be reliable [16]. However, combining multiple sound repetition detections, neuro-imaging methods could be used to probe converging cognitive functions for coma patient [17]. The present analysis that MMN should be replaced by the predictive maps obtained from standard and deviant sounds (Fig. 4). The two classes of sounds are neither redundant nor independent and we used different features to generate classification maps.

### 3.2 Predictive methods

In the past decade, statistical methods based on classification such as logistic regression analysis and single trial topographical analysis [18], Gaussian mixture model estimators [19] were used to classify various brain states, coma, but also performance during therapeutic hypothermia after re-warming to normal body temperature (normothermia) and within 24-48 hours from coma onset [9].

In the present study, we analyzed a single data set that lead to two classification maps. We proposed that taken the minimum of deceased probabilities (relation 20) among several classifiers could be chosen as a safe predictor of the coma outcome. However, collecting multiple data within few hours or days could reveal changes that could have a higher predictive value. It has already been noticed that the progression of auditory discrimination over the first two days of coma is highly predictive for exit from coma [18]. The present method could generalize to account for these time dependent data set.

Finally, we have shown here that modern statistical approaches allow to convert a classification of survivors and deceased patients map into a predictive survival tool. These maps can be refined and upgraded by adding new cases and thus increase the performance of the probabilistic classifier. This approach could be applied to other predictive situations and generalize to other coma types.

## 4 Material and methods

### 4.1 Data acquisition

We collected the data under the auditive mismatch negativity (MMN) protocol CAPACITY, between January 2013 and January 2016. It consists in a 20 minutes EEG session where the patient is subjected to an auditive stimulus each second. Stimuli are divided into two types: standard and deviant. Standard ones consist in a 20ms long sound at 500Hz while deviant stimuli are 30 ms long at 1000Hz. Both signal power is 65dB. Every second, one of the two stimuli is randomly selected with a probability 0.9 for standard ones and 0.1 for deviant ones.

Signals were recorded in 30 patients between 2 and 5 days after cardiac arrest, regardless of sedation. Recordings were performed at electrode sites Fz, Cz, C3, C4, T3, T4 of the International 10-20 system for 20 minutes. Data were obtained following a standard procedure of regular repetitive auditory stimulations (80 dB intensity, 75ms duration and 800Hz frequency) corresponding to the standard curve of the MMN. These data were exported in the European Data Format (EDF), which is a simple and flexible format for storage of multichannel biological and physical signals, then anonymized through a specific software we designed. We focused first on one central electrode Cz and used different time windows (first few hundreds milliseconds) to study the signal fluctuation and correlation between these time windows over 20 minutes of recording.

In the present approach a MMN is a negative peak obtained in the difference between deviant and standard response occuring in the time interval 150-300ms, following stimulation. We collected the periodic responses and calibrated all of them on the initial stimuli, so that no shift was introduced during averaging the responses.

### 4.2 Data pre-processing

We first focus on the evoked signal, corresponding to responses to standard periodic auditory stimuli, repeated every 1s (Fig. 1A green dot). We isolated the CZ-electrode for the evoked response concerning non-deviant stimulations. We filtered the time series *X*(*t*) using a Butterworth bandpass filter (*n* = 4) in the frequency range 0.5-50Hz (Fig. 1B) and obtain the output *X*_*f*_ (*t*). Finally, we average the signal in the time interval [0 − 1]*s*, ensuring that auditory stimuli were produced at time *t* = *nT* (T=1s) leading to the response

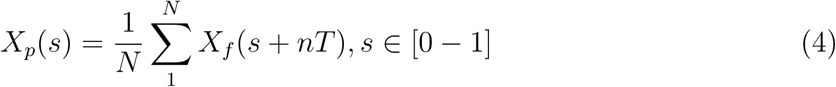

where *N* is the number of periods (typically of the order 10^3^). This preliminary procedure therefore allows to obtain an averaged response *X*_*p*_ that highlights any possible deterministic feature present in the response (FIG 1C).

### 4.3 EEG signal processing and identification of main variables

For the analysis of standard stimulation, we divided the 20 minutes recording into n parts (n=2 or 3) to compute temporal correlations.

#### 4.3.1 Analysis of standard stimuli

We define now the main parameters we extracted to study the response to standard stimulations.

1. We compute the variance of the signal in the time interval [20 − 320]*ms*:

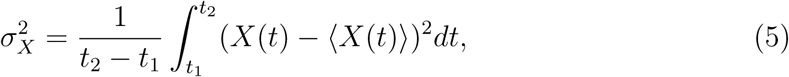

where *t*_2_ = 320*ms* and *t*_1_ = 20*ms*. This time interval corresponds to time scales of the neural networks involved in cognitive tasks.
2. We divide the acquisition time of 20 minutes into n equal parts. For n=2, we get [1 − 10]*min* and [10 − 20]*min*. We average the signals on each of these periods to obtain two responses *X*(*t*) and *Y* (*t*) in the interval [0, 1]*s* (step 1). We computed the time correlation in [20, 320]*ms* of these two signals:

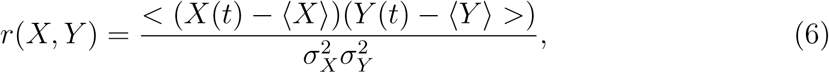

where < . > Represents this time average and m (X) is the average of the variable X.

We conclude that we use two parameter to represent the state of a patient: the variance *s*_*X*_, computed over the entire sample of 20 minutes and the correlation index *r*(*X, Y*), computed in eq. 6.

### 4.4 Analysis of responses to deviant stimuli

Deviant stimuli are 1/10th of the entire responses. To define the signal of interest, we sum the signals from electrodes CZ, C3, C4 and FZ. We then filtered the resulting signal *X*_*d*_ using a lowpass Butterworth filter (n=2) with a cut off frequency at 10Hz (Fig. 1). Finally, we isolated responses on a window of 500*ms* and computed an average responses

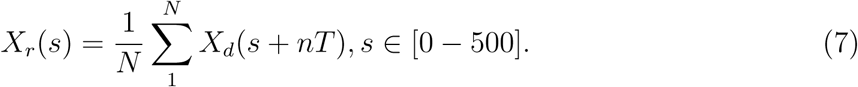

The smooth signal is shown in Fig. 1C and we extracted two features:

1. The number *N*_*E*_ of local extrema (minima and maxima) in the response attained at points *e*_*i*_.
2. The oscillation

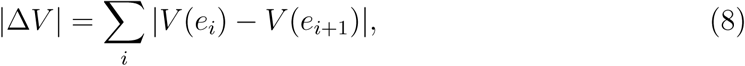

which is the sum of the absolute value of the difference between two consecutive extrema of the average evoked response (see Fig. 3).

### 4.5 Two-dimensional representation

Based on the parameters we extracted in the previous subsections 4.3.1-4.4, we generated two-dimensional maps: for the map associated to non-deviant stimuli, each patient has coordinates *P* = (*σ*_*X*_, *r*(*X, Y*)), while for deviant stimuli, we use the coordinates *P* = (*N*_*E*_, |Δ*V*|). In various plot, we normalized the coordinates in a population (***X***_1_,..***X***_*n*_) by:

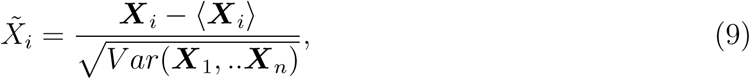

where ⟨***X***_*i*_⟩ is the average over the points ***X***_*i*_ and Var is the variance.

We map all points for all patients as shown in fig. 4, with the color code that deceased (resp. survived) patient are shown in blue (resp. red). We note that survived patients form cluster that will be the basis of the classification and prediction described below.

### 4.6 Classifiers to compute the Survival probability

To use the maps defined in subsection 4.5 as predictive tool, we use two statistical classifiers. Indeed, we wish to assign a survival probability to any point present or that could be added on the map based on the ensemble of previous data points already classified. Using the assumption that statistics associated to patients (features) are independent one from each other, we used a Bayesian classification.

#### 4.6.1 SVM Classification

To classify the date, we use the standard SVM algorithm [20] which finds the hyperplane that best separates the two classes (deceased vs survived). The hyperplane maximizes the distance between itself and the closest points of each class, while all element of each class is located on each of the two sides [21]. If no such hyperplane is found, which is the case here, the dimension of the space where the data are embedded should be increased, a procedure known as kernelling [22]. In this higher dimensional space, the classes are well separated by a higher dimensional hyperplane. If the two classes are still not well separated, a penalty is inflicted for every misclassified data point [23]. Here, the kernel is the Radial Basis Function *K*(*x, x*^*′*^) = exp(−*γ*‖*x* − *x*^*′*^‖^2^), with *γ* = 1 and a penalty coefficient *C* = 10. We implemented the SVM using the Scikit Learn module [24, 25].

#### 4.6.2 Gaussian estimator

In case of a Gaussian estimator, we estimated the mean and the covariance matrix for the survival and deceased classes. The probability of each class is computed empirically using the maximum likelihood estimator (see appendix). We recall for an ensemble of n data 𝒮_*n*_ = (***x***_1_,..***x***_*n*_) that are separated into two classes, *C*_1_ and *C*_2_, the probability that a patient ***X*** belongs to one class, conditioned on the ensemble 𝒮_*n*_:

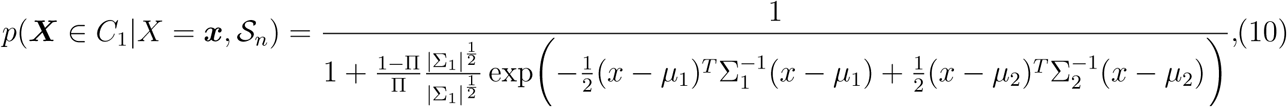

where (*µ*_*i*_, Σ_*i*_)_*i*=1,2_ are the mean and variance computed from each class *C*_1_ and *C*_2_ from 𝒮_*n*_. We used the fraction 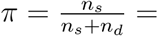, for the number of survival *n*_*s*_ and deceased patients *n*_*d*_. Formula 10 is derived in the appendix.

#### 4.6.3 K-Nearest neighbors classifier

To classify the standard stimuli, we use the K-Nearest neighbors classifier. We computed the ratio for the probability of belonging to a class. For a given point *X*, the probability to belong to class *C*_1_(“Survival”) given the distribution of point ***x*** is computed empirically as the number of surviving neighbors out of a total of K.

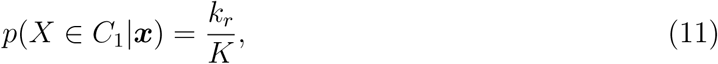

where *k*_*r*_ is the number of neighbors that belongs to the class “Survival” among K closest points.

#### Weighted K-Nearest Neighbors

To classify deviant stimuli, we use a variant of the K-neighbors method by adding distance-relative weights to the points inside the dataset. The two classes labeled “ Deceased” and “Survival” are defined as *C*_1_ and *C*_2_ respectively. The ensemble of points 𝒮_*n*_in dimension 2 are given by the coordinates ***x***_*n*_ = (*N*_*E,n*_, Δ*V*_*n*_), extracted in subsection 4.4. To compute the classification probability, we define *K* −nearest neighborhood 𝒩_*K*_(***x***) for the point ***x*** as the *K* shortest points from ***x***, computed from the Euclidean distance (between two points ***x***_*n*_, ***x***_*m*_),

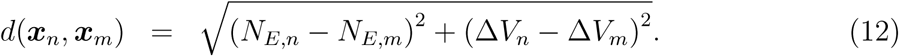

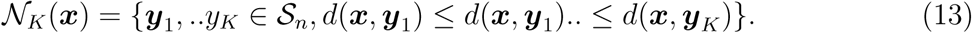

Finally, the conditional probability is

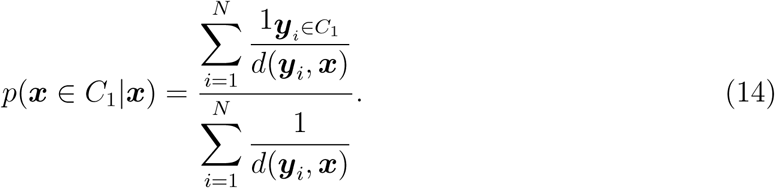

#### 4.6.4 Cross-validation

We used a cross-validation approach to validate the classification algorithm: we excluded a patient at a time and computed the survival probability, based on the remaining elements in the data basis [26]. We then computed the survival probability using the three classifiers K-neighbors, SVM and the Gaussian estimators and compared the result to the true result. We followed the protocole: 1-a patient *P*_*i*_, *i* = 1..*N* is selected inside the data basis. 2-we trained the classification algorithm on the database of all patient {*P*_*k*_, *k* = 1*..N*} − *P*_*i*_. We evaluate the prediction of the model on the excluded patient, leading to a score *s*_*i*_. We recall that *s*_*i*_ = 1 if the prediction is correct, otherwise *s*_*i*_ = 0. We then replace the patient *P*_*i*_ inside the data base and reiterate procedure until each patient has been exactly excluded once. The final score of the model is computed as

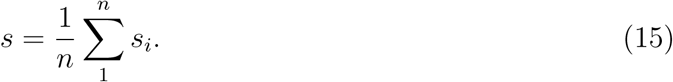

Finally, the confusion matrix defined as

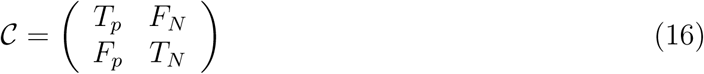

for the true positive *T*_*p*_ (number of patient that have survived and classified correctly), true negative *T*_*n*_ (number of patient that have deceased classified correctly) and false positive *F*_*p*_(number of patient that have survived and classified incorrectly) and false negative *F*_*n*_ (number of patient that have deceased and classified incorrectly). The success of classification is given by

1. the Accuracy (rate of patients correctly classified):

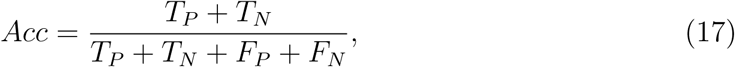
2. the Sensibility is the rate of survival patient correctly classified, defined empirically by

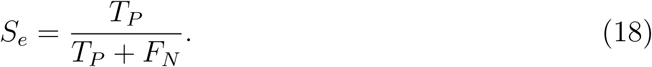
3. The Specificity is the rate of deceased patient correctly classified

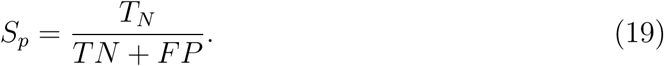

This cross-validation shows that our classifier on deviant or non-deviant patient gave very accurate prediction with a worth rate > 90% success (for Gaussian) and not mistake 100% using k-neighbors for *k* = 3 and 97% success for *n* = 4, see table 2.

#### 4.6.5 Combined probability

We propose to use for the predictive decisional probability *p*_*dec*_ the minimum of the ones estimated for the deviant (relation 14) and non-deviant (relation 11) classifications. For a patient of coordinate ***x*** in each map has survival probability:

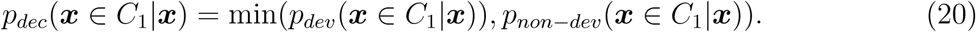

#### 4.6.6 Iteration and changing k-neighbors k

The approach developed here is iterative and any new additional case enrich the data base and the classifications maps. For the K-neighbors approach, adding a point do not require any changes in the computation, although we expect that the number of neighbors will enter into the computation could diminish as the number of cases added in the map increases. For the Gaussian classification, the mean and the variance are recomputed following each new cases.

## Appendix

### 5.3 Computing the Apriori estimators

We compute here the apriori Gaussian empirical estimators. In the framework of two classes for survival and deceased patients that we consider to be normally multidimensional distributed, we will first derive the estimator for a point ***x*** to be classified, based on the mean and variance, that we relate to the sample of the database of surviving patients. The computations use the Bayes’rule and we assume that each class can be distinguished by its mean and covariance matrix, which should a priori be different. The variable *y* represents the classification to one of the class, while *x* represents the position in the phase space. We assume the following apriori probability:

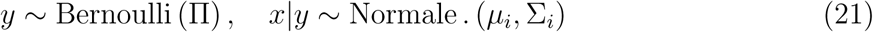

The parameter to be estimated are *θ* = (Π, *µ*_0_, *µ*_1_, Σ_0_, Σ_1_). The data base 𝒮_*n*_ is of size *n*. The log-likelihood estimator is

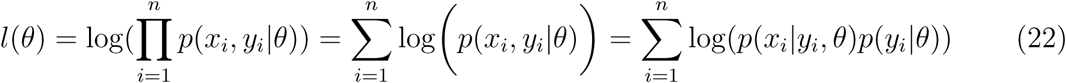

Splitting the sum with respect to the two classes represented by *y* = 1 for (*p*(*y*|*θ*) = Π) and *y* = 0 (*p*(*y*|*θ*) = 1 − Π), we get:

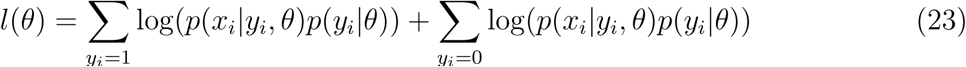

Then:

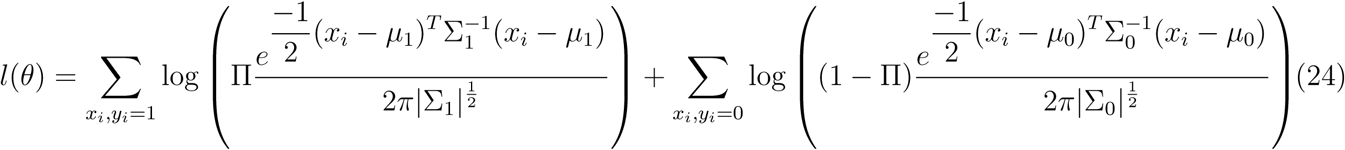

The total number of points in class 1 is

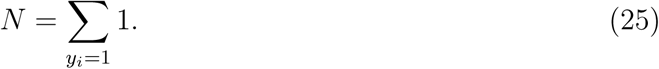

We finally get:

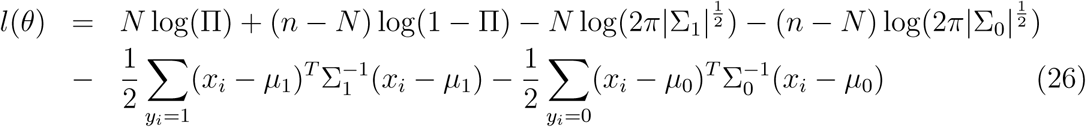

The value of the different parameter are obtained at the extremum of the estimators. Thus,

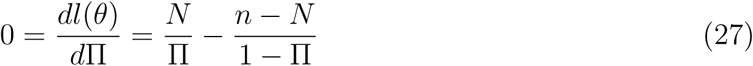

leading to the empirical estimator

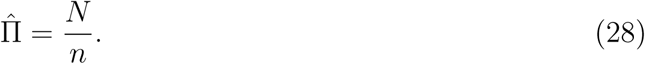

Differentiating with the mean, we get:

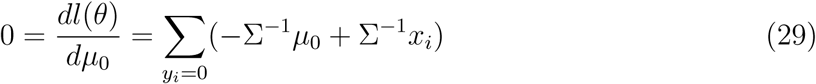

Thus

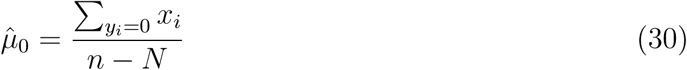

Similarly,

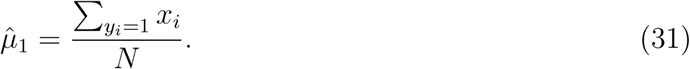

Finally,

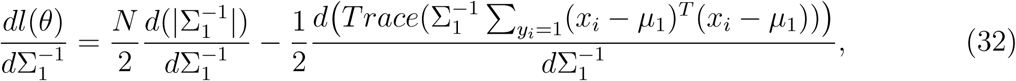

and we recall that 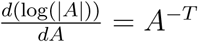 et 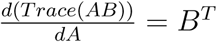, indeed:

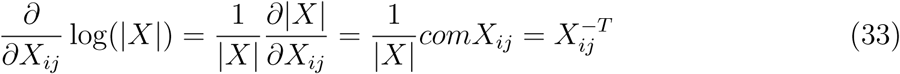

Using that (*comA*)^*T*^ = |*A*|*A*^*-*1^ for an invertible matrix, we get

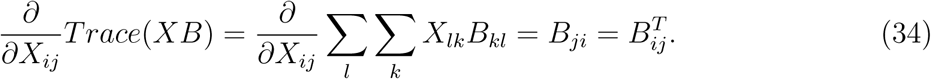

Then,

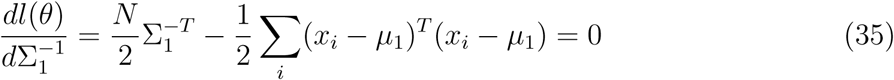

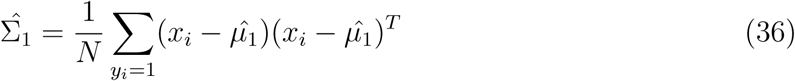

Similarly

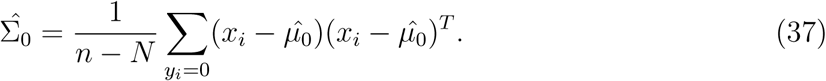

We conclude with the final apriori probability:

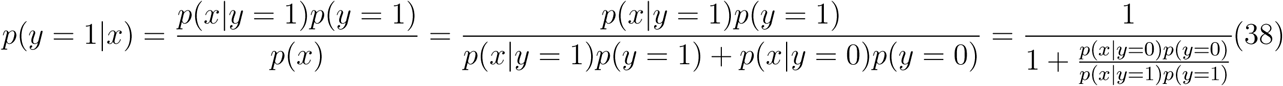

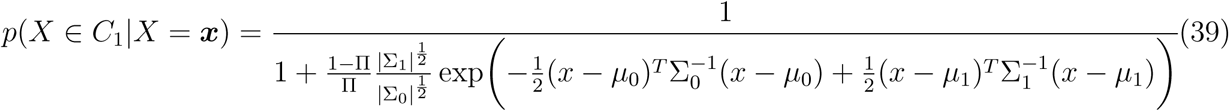

We use relation 39 to estimate the probability for a point ***x*** to below to a given class, after the parameters are estimated from the ensemble of data.

## References

[1] N. André-Obadia, J. Zyss, M. Gavaret, J.-P. Lefaucheur, E. Azabou, S. Boulogne, J.-M. Guerit, A. McGonigal, P. Merle, V. Mutschler, et al., “Recommendations for the use of electroencephalography and evoked potentials in comatose patients,” Neurophysiologie Clinique, 2018.

[2] U. Schall, “Is it time to move mismatch negativity into the clinic?,” Biological psychology, vol. 116, pp. 41–46, 2016.

[3] C. C. Duncan, R. J. Barry, J. F. Connolly, C. Fischer, P. T. Michie, R. Näätänen, J. Polich, I. Reinvang, and C. Van Petten, “Event-related potentials in clinical research: guidelines for eliciting, recording, and quantifying mismatch negativity, p300, and n400,” Clinical Neurophysiology, vol. 120, no. 11, pp. 1883–1908, 2009.

[4] C. Fischer, D. Morlet, P. Bouchet, J. Luaute, C. Jourdan, and F. Salord, “Mismatch negativity and late auditory evoked potentials in comatose patients,” Clinical neurophysiology, vol. 110, no. 9, pp. 1601–1610, 1999.

[5] D. Gabriel, E. Muzard, J. Henriques, C. Mignot, L. Pazart, N. Andr-Obadia, J.-P. Ortega, and T. Moulin, “Replicability and impact of statistics in the detection of neural responses of consciousness,” Brain, vol. 139, no. 6, p. e30, 2016.

[6] M. De Lucia and A. Tzovara, “Reply: Replicability and impact of statistics in the detection of neural responses of consciousness,” Brain, vol. 139, no. 6, pp. e32–e32, 2016.

[7] L. Naccache, L. Puybasset, R. Gaillard, E. Serve, and J.-C. Willer, “Auditory mismatch negativity is a good predictor of awakening in comatose patients: a fast and reliable procedure,” Clinical Neurophysiology, vol. 116, no. 4, pp. 988–989, 2005.

[8] J. Daltrozzo, N. Wioland, V. Mutschler, and B. Kotchoubey, “Predicting coma and other low responsive patients outcome using event-related brain potentials: a meta-analysis,” Clinical Neurophysiology, vol. 118, no. 3, pp. 606–614, 2007.

[9] M. De Lucia and A. Tzovara, “Decoding auditory eeg responses in healthy and clinical populations: a comparative study,” Journal of neuroscience methods, vol. 250, pp. 106–113, 2015.

[10] L. Naccache, J.-R. King, J. Sitt, D. Engemann, I. El Karoui, B. Rohaut, F. Faugeras, S. Chennu, M. Strauss, T. Bekinschtein, et al., “Neural detection of complex sound sequences or of statistical regularities in the absence of consciousness?,” Brain, vol. 138, no. 12, pp. e395–e395, 2015.

[11] E. Juan, M. De Lucia, A. Tzovara, V. Beaud, M. Oddo, S. Clarke, and A. O. Rossetti, “Prediction of cognitive outcome based on the progression of auditory discrimination during coma,” Resuscitation, vol. 106, pp. 89–95, 2016.

[12] J. Hofmeijer, M. C. Tjepkema-Cloostermans, and M. J. van Putten, “Burst-suppression with identical bursts: a distinct eeg pattern with poor outcome in postanoxic coma,” Clinical neurophysiology, vol. 125, no. 5, pp. 947–954, 2014.

[13] J. Hofmeijer and M. J. A. M. van Putten, “Eeg in postanoxic coma: prognostic and diagnostic value,” Clinical neurophysiology, vol. 127, no. 4, pp. 2047–2055, 2016.

[14] J. Friedman, T. Hastie, and R. Tibshirani, The elements of statistical learning, vol. 1. Springer series in statistics New York, NY, USA:, 2001.

[15] J. Luaute, C. Fischer, p. Adeleine, D. Morlet, L. Tell, and D. Boisson, “Late auditory and event-related potentials can be useful to predict good functional outcome after coma,” Archives of physical medicine and rehabilitation, vol. 86, no. 5, pp. 917–923, 2005.

[16] E. Azabou, B. Rohaut, R. Porcher, N. Heming, S. Kandelman, J. Allary, G. Moneger, F. Faugeras, J. Sitt, D. Annane, et al., “Mismatch negativity to predict subsequent awakening in deeply sedated critically ill patients,” British journal of anaesthesia, vol. 121, no. 6, pp. 1290–1297, 2018.

[17] C. Sergent, F. Faugeras, B. Rohaut, F. Perrin, M. Valente, C. Tallon-Baudry, L. Cohen, and L. Naccache, “Multidimensional cognitive evaluation of patients with disorders of consciousness using eeg: a proof of concept study,” NeuroImage: Clinical, vol. 13, pp. 455–469, 2017.

[18] A. Tzovara, A. O. Rossetti, L. Spierer, J. Grivel, M. M. Murray, M. Oddo, and M. De Lucia, “Progression of auditory discrimination based on neural decoding predicts awakening from coma,” Brain, vol. 136, no. 1, pp. 81–89, 2013.

[19] M. De Lucia, A. Tzovara, F. Bernasconi, L. Spierer, and M. M. Murray, “Auditory perceptual decision-making based on semantic categorization of environmental sounds,” Neuroimage, vol. 60, no. 3, pp. 1704–1715, 2012.

[20] V. Cherkassky and F. Mulier, Learning from data: Concepts, theory, and methods. Wiley New York, 1998.

[21] L. G. Valiant, “A theory of the learnable,” Communications of the ACM, vol. 27, no. 11, pp. 1134–1142, 1984.

[22] M. A. Aizerman, “Theoretical foundations of the potential function method in pattern recognition learning,” Automation and remote control, vol. 25, pp. 821–837, 1964.

[23] C. Cortes and V. Vapnik, “Support-vector networks,” Machine learning, vol. 20, no. 3, pp. 273–297, 1995.

[24] F. Pedregosa, G. Varoquaux, A. Gramfort, V. Michel, B. Thirion, O. Grisel, M. Blondel, p. Prettenhofer, R. Weiss, V. Dubourg, J. Vanderplas, A. Passos, D. Cournapeau, M. Brucher, M. Perrot, and E. Duchesnay, “Scikit-learn: Machine learning in Python,” Journal of Machine Learning Research, vol. 12, pp. 2825–2830, 2011.

[25] L. Buitinck, G. Louppe, M. Blondel, F. Pedregosa, A. Mueller, O. Grisel, V. Niculae, p. Prettenhofer, A. Gramfort, J. Grobler, R. Layton, J. VanderPlas, A. Joly, B. Holt, and G. Varoquaux, “API design for machine learning software: experiences from the scikit-learn project,” in ECML PKDD Workshop: Languages for Data Mining and Machine Learning, pp. 108–122, 2013.

[26] R. Kohavi et al., “A study of cross-validation and bootstrap for accuracy estimation and model selection,” in Ijcai, vol. 14, pp. 1137–1145, Montreal, Canada, 1995.

